# Novel polymeric fluoropyrimidine CF10 demonstrates superior therapeutic index and survival advantage in patient-derived models of 5-fluorouracil-refractory colorectal cancer

**DOI:** 10.64898/2026.04.05.716582

**Authors:** Naresh Sah, Tasmin R. Omy, Subash Kairamkonda, Ganesh Acharya, Hithardha Palle, Pamela Luna, Chinnadurai Mani, William Gmeiner, Naga Cheedella, Mark B. Reedy, Komaraiah Palle

## Abstract

**Background:** Fluoropyrimidines, specifically 5-fluorouracil (5-FU), remain the cornerstone of colorectal cancer (CRC) therapy. However, intrinsic and acquired resistance, alongside dose-limiting systemic toxicities, often result in treatment failure and disease relapse. There is a pressing clinical need for next-generation fluoropyrimidines that can retain the antitumor activity in 5-FU-refractory CRC models while maintaining a favorable safety profile.

**Methods:** We evaluated the antitumor efficacy of CF10, a novel polymeric fluoropyrimidine designed for the sustained delivery of FdUMP, against equimolar 5-FU. We utilized a diverse panel of six patient-derived CRC organoid (PDO) models to assess 3D growth inhibition under both normoxic (∼20% O_2_) and physioxic (5% O_2_) conditions. Mechanisms of action were investigated via γH2AX signaling (DNA damage), Annexin V/PI flow cytometry (death kinetics), and ALDEFLUOR™ assays (stem-like populations). Functional suppression of metastasis-associated phenotypes was evaluated using 3D Matrigel invasion assays. Finally, the therapeutic index and overall survival were validated *in vivo* using two independent patient-cell-derived xenograft (PCDX) models (TX-CC-199 and TX-CC-201).

**Results:** CF10 demonstrated significantly greater suppression of organoid growth compared to equimolar 5-FU across all patient-derived lines, regardless of morphological heterogeneity or oxygen tension. In 3D invasion assays, CF10 achieved superior anti-invasive activity even at a 10-fold lower molar dose than 5-FU. This functional advantage was mirrored by a marked depletion of the ALDH-high stem-like subpopulation, which was largely recalcitrant to 5-FU. Mechanistically, CF10 induced intensified replication stress, DNA damage and repair signaling (γH2AX, Top1cc/pRPA32, FANCD2), and pushed the CRC to irreversible/terminal, PI-positive death states. *In vivo*, CF10 treatment resulted in profound tumor growth inhibition and a robust survival advantage in two patient cell-derived xenograft (PCDX) models (Log-rank P<0.01) without inducing systemic weight loss or noticeable toxicity.

**Conclusions:** By integrating 3D patient-derived modeling with *in vivo* validation, we demonstrate that CF10 effectively overcomes the biological and pharmacological limitations of 5-FU. CF10 targets the aggressive, invasive, and stem-like subpopulations of CRC that drive clinical relapses. These findings provide a compelling translational rationale for the clinical development of CF10 as a superior alternative to standard fluoropyrimidines in both treatment-naive and refractory CRC.

**Significance Statement:** Despite the foundational role of 5-fluorouracil (5-FU) in colorectal cancer (CRC) therapy, resistance and systemic toxicity remain major barriers to curative outcomes. This study identifies CF10, a novel polymeric fluoropyrimidine, as a superior alternative that overcomes 5-FU resistance in biologically diverse patient-derived organoids and xenograft models.

Crucially, CF10 demonstrates a unique capacity to suppress the invasive, aldehyde dehydrogenase (ALDH)-high stem-like subpopulations that likely survive standard chemotherapy (5-FU) by maintaining efficacy under physiological oxygen levels and providing a significant survival advantage *in vivo* with improved tolerability. CF10 represents a promising translational candidate for the treatment of both treatment-naive and refractory CRC.

## Introduction

Colorectal cancer (CRC) remains the second leading cause of cancer-related mortality worldwide and is projected to claim over 500,000 lives in 2026 alone, despite significant advancements in screening and systemic intervention.^1,2^ Fluoropyrimidines, specifically 5-fluorouracil (5-FU), remain the cornerstone of CRC treatment across both adjuvant and metastatic settings.^3,4^ However, the clinical utility of 5-FU is frequently compromised by intrinsic and acquired resistance, as well as dose-limiting toxicities that preclude durable responses and lead to refractory (incomplete responses and relapse) disease.

In recent years, the therapeutic landscape for fluoropyrimidine-refractory CRC has shifted toward biomarker-driven, regimen-level strategies rather than optimizations of the fluoropyrimidine backbone itself. These include (i) targeted combinations for molecular subsets, such as encorafenib/cetuximab for *BRAF*V600E mutations and emerging *KRAS*G12C inhibitors^5–7^; (ii) immune checkpoint blockade, such as pembrolizumab, for MSI-H/dMMR disease^8–10^; and (iii) the use of agents that maintain activity post-5-FU exposure, including trifluridine/tipiracil (TAS-102) combined with bevacizumab^11^ and small-molecule inhibitors like fruquintinib^12^ Together, these advancements highlight two persistent unmet needs: clinical progress increasingly relies on alternative therapies that bypass 5-FU resistance mechanisms rather than improving the fluoropyrimidine backbone itself, and these advanced therapies must be evaluated in models that better recapitulate patient tumor biology. These needs remain clinically relevant because screening and molecularly/biomarker-guided advances do not eliminate the large subset of patients who present with advanced or refractory CRC, or harbor tumors that are stubbornly resistant to existing systemic regimens. To improve the fluoropyrimidine backbone itself, we have developed CF10, a novel polymeric fluoropyrimidine designed for the sustained delivery of the thymidylate synthase (TS)-directed antimetabolite, 5-fluoro-2’-deoxyuridine-5’-monophosphate (FdUMP).^13–15^ CF10 is engineered to intensify tumor-selective cytotoxicity while improving systemic tolerability.

Traditional preclinical evaluation often relies on long-term established 2D cell lines, which fail to capture the inter-patient heterogeneity and 3D architecture that govern therapeutic response.^16,17^ To address this failure, patient-derived organoids (PDOs) have emerged as a robust bridge between 2D cultures and *in vivo* models, preserving key tumor-intrinsic features and enabling direct pharmacological comparisons.^18–20^ Critically, drug sensitivity is modulated by the physiological constraints inherent to 3D growth, such as physioxia, cell-cell interactions and nutrient gradients.^21,22^ This is particularly relevant in CRC, because oxygen availability (physioxia vs. normoxia) and microenvironmental factors significantly influence DNA damage responses and HIF-driven resistance pathways.^23–26^ Additionally, aggressive CRC is defined by its capacity for invasion and persistence, often linked to epithelial-mesenchymal plasticity (EMP) and the acquisition of stem-like properties^27–29^ In particular, the aldehyde dehydrogenase high (ALDH-high) population serves as a critical functional marker for these tumor-initiating cell states.^30,31^

In this study, we provide a comprehensive evaluation of CF10 antitumor activity compared to equimolar 5-FU. We utilize multiple patient-derived primary CRC cultures to assess 3D growth inhibition under both normoxic and physioxic (low oxygen) conditions. We further delineate the mechanisms of efficacy by quantifying DNA-damage signaling, cell-death kinetics, and 3D invasive outgrowth. To bridge the gap to whole-organism pharmacology, we extended these findings into patient-cell-derived xenograft (PCDX) models. By integrating 3D modeling, functional stemness assays, and *in vivo* efficacy, we demonstrate that CF10 offers a clinically meaningful advantage over 5-FU in eradicating aggressive tumor cell states.

## Material and methods

### Patient-derived colorectal cancer (CRC) organoid culture

Six primary CRC lines (TX-CC-201, TX-CC-199, TX-CC-286, TX-CC-096h2, TX-CC-208, TX-CC-206) were received from the TTUHSC cancer center. Cells were maintained in Iscove’s Modified Dulbecco’s Medium (IMDM; Gibco #12440-053) supplemented with 20% fetal bovine serum (FBS; heat-inactivated), 1% insulin-transferrin-selenium (ITS; Gibco #41400-045), 100 U/mL penicillin, and 100 μg/mL streptomycin at 37°C, 5% CO₂ (normoxia). Where indicated, cultures were adapted and maintained in physioxia (TX-CC-096h2, TX-CC-286, and TX-CC-199) using a tri-gas incubator, Whitley H35 Hypoxystation (Don Whitley Scientific Ltd., Bingley, UK), at 5% O₂ (unless otherwise noted in figure legends). Cells were routinely screened for Mycoplasma (Mycoplasma PCR Detection Kit; Abcam #AB289834) and used at ≤10 passages after thawing for all experiments.

### Drugs and Reagents

CF10 was prepared as previously described by our group and supplied as a sterile aqueous stock (store at −80 °C).^14,15,32^ 5-Fluorouracil (5-FU; (50 mg/mL; NDC 63323-117-00, Fresenius Kabi USA, LLC, Lake Zurich, IL) was dissolved in sterile saline to prepare working stocks, which were sterile filtered through a 0.22 μm PVDF filter before use in cell culture. Matrigel® Growth Factor-Reduced (Catalog # 3500-096-K; R&D Systems, a Bio-Techne brand, Minneapolis, MN, USA) was used for 3D invasion assays. Unless stated otherwise, all other reagents were analytical grade.

### 3D patient-derived organoid drug-response assay (imaging-based stratification)

Patient-derived colorectal cancer (CRC) cultures (TX-CC lines, as indicated) were expanded under standard conditions before 3D assays. For organoid formation, cultures were dissociated to single cells using trypsin, counted, and seeded at ∼1 × 10⁴ cells per well (technical triplicates) in ultra-low attachment 6-well plates (Cat. #3471, Corning) in IMDM-based medium. After waiting 24-48 hrs to allow aggregation, organoids were treated with vehicle, CF10, or 5-FU at various concentrations (as provided in the figure legends). Where indicated, assays were conducted under physioxic conditions (5% O₂) using a Whitley H35 Hypoxystation (Don Whitley Scientific Ltd., Bingley, UK), with matched plates maintained under normoxia.

For the primary CRC line TX-CC-096h2, which formed continuous sheet/tubule-like organoid networks under 3D culture rather than discrete organoids, organoid growth was quantified as projected organoid area per field instead of organoid size. For the round, discrete organoid-type organoids, growth was quantified as the size (µm) and the number of organoids rather than the projected area. At the endpoint (after around 12-14 days of treatment), brightfield images were acquired on a Nikon Eclipse Ts2 inverted microscope (Nikon Instruments Inc., Melville, NY, USA) equipped with a 4× objective and a digital camera, using identical exposure and acquisition settings across all conditions. For each well, five non-overlapping fields (center and four quadrants) were imaged, yielding 5 fields per condition per plate; experiments were repeated on three independent plates (n = 3 biological replicates). Brightfield TIFF images were batch-quantified in Fiji (ImageJ; Fiji v2.16.0, ImageJ v1.54p; Java 21.0.7, 64-bit) using a custom macro (8-bit conversion, rolling-ball background subtraction, Otsu thresholding, binary masking, and area measurement); the full macro is provided as Supplementary Macro S1.

### 3D Matrigel organoid invasion

The invasive potential of 3D cultures was assessed using the Cultrex® 3-D Organoid BME Cell Invasion Assay (Cat. #3500-096-K; R&D Systems/Bio-Techne, Minneapolis, MN, USA) with minor modifications to the manufacturer’s protocol. Briefly, cells were seeded into low-attachment, suspension culture 96-well plates (Cat # 655185, CellStar, Greiner Bio-One, Monroe, NC, USA) at 1-2 × 10^3 cells/well in Organoid medium (IMDM supplemented with 20% FBS and 0.5% ITS) and allowed to form Organoids for 48 h. Preformed organoids were treated with complete medium containing equimolar CF10, 5-FU, or vehicle control, and invasion was monitored for up to 7 days.

Brightfield images were acquired on Days 0, 3, and 7 (40X) using identical acquisition settings across conditions. Invasion was quantified in a blinded manner in Fiji/ImageJ using a standardized segmentation workflow to measure total organoid footprint (core + outgrowth) and core organoid area; invasion area was calculated as Total Area − Core Area. For longitudinal analyses, values were normalized to the mean vehicle control for the corresponding day. At least 3 organoids per condition were quantified per experiment; data are presented as mean ± SEM.

### ALDEFLUOR™ assay in patient-derived CRC organoids

The ALDH-high cell population was quantified using the ALDEFLUOR™ Assay Kit (STEMCELL Technologies; Cat. #01700) according to the manufacturer’s instructions with minor modifications for patient-derived 3D CRC cultures. Briefly, TX-CC-096h2 and TX-CC-199 cultures were treated for 72 h with vehicle, equimolar 5-FU, or CF10, then harvested for flow-cytometric analysis. Organoid cultures were collected, washed with PBS, and dissociated into single cells. Dissociated cells were counted and resuspended in ALDEFLUOR assay buffer at approximately 1 × 10⁶ cells/mL. For staining, cells were incubated with the ALDEFLUOR fluorescent substrate (BAAA) prepared in assay buffer. For each condition, a matched negative-control aliquot was treated with the ALDH inhibitor diethylaminobenzaldehyde (DEAB; 50 mmol/L) to define background fluorescence and establish the ALDH⁻ gate. After incubation under the kit-recommended conditions, cells were washed and maintained on ice until acquisition.

Samples were acquired on a Beckman Coulter flow cytometer and analyzed using Kaluza software (Beckman Coulter). A standard gating strategy was applied: debris was excluded using FSC/SSC, gating prior to ALDH analysis. The ALDH⁺ gate was set using the corresponding DEAB control and applied consistently across treatment groups within each experiment. Experiments were performed in technical triplicate per condition and repeated across three independent experiments. Results are reported as percentage ALDH⁺ cells per condition with representative gating plots and summary quantification.

### Flow cytometry for apoptosis (Annexin V/PI)

Apoptosis was assessed 72 hrs after treatment using an Annexin V–FITC/propidium iodide (PI) apoptosis detection kit (Invitrogen/eBioscience; Cat. #BMS500FI-100). Cultures were harvested and dissociated to single cells using trypsin, washed with cold PBS, and resuspended in 1× Annexin V binding buffer. Cells were incubated with Annexin V–FITC for 15 min at room temperature, protected from light, followed by the addition of PI immediately before acquisition. Samples were acquired on a Beckman Coulter flow cytometer, and data were analyzed using Kaluza (Beckman Coulter) software. Debris and doublets were excluded by FSC/SSC and area–height gating. Cell death states were quantified as live (Annexin V⁻/PI⁻), early apoptotic (Annexin V⁺/PI⁻), and late apoptotic/secondary necrotic (Annexin V⁺/PI⁺). Quadrant gates were set up using unstained and single-stained controls. A minimum of 30,000 events per sample were collected. Experiments were performed in 3 independent triplicates.

### Western blotting

Cells were placed on ice, washed twice with ice-cold PBS, and lysed in ice-cold RIPA buffer supplemented with protease and phosphatase inhibitors (cOmplete™ and PhosSTOP™, Roche) as described previously.^33,34^ Lysates were clarified by centrifugation at 4 °C, and the supernatants were collected. Protein concentration was determined using a BCA assay (Thermo Fisher Scientific). Equal amounts of protein (20–40 µg) were mixed with Laemmli sample buffer, denatured by heating, and resolved on 10% SDS–PAGE gels. Proteins were transferred to 0.45 µm PVDF membranes using Trans-Blot Turbo Transfer System (Bio-Rad; catalog #1704150) and Transfer Buffer (Bio-Rad; catalog #10026938). Membranes were blocked in 5% non-fat milk in TBST (Tris-buffered saline, 0.1% Tween-20) and incubated overnight at 4 °C with primary antibodies (typically 1:1,000) against γH2AX (γH2AX; Ser139, Cell Signaling Technology, Danvers, MA; Cat. #2577), FANCD2 (FI17; Santa Cruz Biotechnology, Cat. #20022) and Vinculin (loading control) (Cell Signaling Technology; Cat #13901S). After three washes in TBST, membranes were incubated with HRP-conjugated secondary antibodies (1:2,000) for 1 hr at room temperature, washed again, and developed using enhanced chemiluminescence (SuperSignal™ West Femto, Thermo Fisher Scientific). Chemiluminescent signals were captured on an autoradiography film in a darkroom.

### Immunofluorescence staining and confocal imaging (intact 3D organoids)

Immunofluorescence (IF) was performed on intact patient-derived CRC organoids to assess DNA damage (γH2AX) and proliferation (Ki67). Organoids were treated as indicated, gently collected to preserve 3D architecture, washed with PBS, and fixed in 4% paraformaldehyde (PFA) for 45 min at room temperature. Fixed organoids were permeabilized with 0.2% Triton X-100 for 10 min and blocked in 3% BSA in PBS. Samples were incubated with primary antibodies against γH2AX-Ser139 (Cell Signaling Technology; Cat. #2577; 1:1000), Top1 (MilliporeSigma; Cat. #MABE1084; 5 μg/mL), pRPA32-S33 (Bethyl Laboratories, Cat. A300-246A; 1:1000), and Ki67 (Cell Signaling Techology; Cat. #9449S; 1:2000) overnight at 4 °C with gentle agitation. After washing, organoids were incubated with Alexa Fluor–conjugated secondary antibodies (1:1,000) for 1 hr at room temperature protected from light, counterstained with DAPI (1 µg/mL), and mounted intact using ProLong™ Gold Antifade (Thermo Fisher Scientific). Immunofluorescence images were acquired on a Nikon Ti-E/A1 confocal microscope and analyzed in ImageJ. Z-stacks were collected using consistent acquisition settings within each experiment. γH2AX foci and Ki67 positivity were quantified using standardized analysis workflows applied uniformly across conditions.

### Patient-derived CRC Model provenance and CDX cohorts’ preparation for treatment studies

Patient-derived colorectal cancer (CRC) models (TX-CC-199 and TX-CC-201) were originally established from fresh colorectal cancer tissue and cryopreserved in the TTUHSC cancer center. Upon receipt, vials were thawed and expanded in vitro to generate sufficient cells for in vivo studies and to optimize xenograft establishment conditions. For CDX generation, expanded cells were counted and viability confirmed before subcutaneous implantation into immunodeficient mice. Two independent CDX cohorts were established from TX-CC-199 and TX-CC-201 and used for randomized treatment studies.

### Cell-derived xenograft (CDX) implantation, randomization, treatment, and monitoring

All animal procedures were conducted under an approved Institutional Animal Care and Use Committee (IACUC) protocol and in accordance with institutional guidelines. Two independent CDX models were established by subcutaneous implantation of patient-derived colorectal cancer (CRC) cells into female nude mice (6–8 weeks): TX-CC-199 (1.5 × 10⁶ cells/mouse) and TX-CC-201 (1.0 × 10⁶ cells/mouse). When tumors reached approximately 100–150 mm³, mice were randomized into vehicle control, 5-FU, or CF10 treatment arms (n = 10 per group, unless otherwise indicated). Drugs were administered intraperitoneally (i.p.) according to the predefined schedule: 5-FU (40 mg/kg, 2×/week) and CF10 (300 mg/kg, 2×/week). Following completion of dosing (through Day 33), animals were monitored without further treatment for survival. Tumor dimensions (length *L* and width *W*) and body weight were recorded twice weekly, and tumor volume was calculated as: 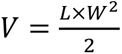. Measurements were performed by an investigator blinded to treatment group whenever feasible. Humane endpoints were predefined and included ulceration, excessive tumor burden (e.g., >1,500 mm³), sustained body-weight loss (>15%), or evidence of moribund condition/distress, consistent with the approved protocol. Primary endpoints included tolerability (body-weight change), and overall survival, with survival analyzed by the Kaplan–Meier method and group comparisons performed using the log-rank (Mantel–Cox) test (as specified in figure legends).

### Statistical analysis

Statistical analyses were performed in GraphPad Prism (v9.5.1). Data are shown as mean ± SEM (unless stated otherwise). For experiments with ≥3 groups, one-way ANOVA with Tukey’s (all pairwise) or Dunnett’s (vs control) multiple-comparisons test was applied; for analyses involving two factors (e.g., treatment × time), two-way ANOVA with multiple-comparisons correction was used (Šidák or Tukey, as indicated). For pairwise comparisons of immunofluorescence intensity measurements, two-tailed Mann–Whitney tests were used. Longitudinal body weight data were analyzed using a mixed-effects model (REML) to accommodate missing values. For in vivo endpoint comparisons (tumor volume and tumor weight), unpaired two-tailed Student’s t-tests were used (with Welch’s correction when variances were unequal). Survival was analyzed by Kaplan–Meier curves and compared using the log-rank (Mantel–Cox) test. All tests were two-sided, and P < 0.05 was considered statistically significant. The details and tests are indicated in the figure legends.

## Results

### CF10 effectively inhibits CRC patient-derived organoid growth compared to 5-FU

Given that 5-FU-based regimens are the cornerstone of first-line CRC treatment, we compared CF10 directly to equimolar 5-FU using multiple PDOs of CRC models^35,36^ since these low-passage primary CRC PDOs provide a clinically relevant platform that preserves tumor-intrinsic heterogeneity.^37,38^ We employed six distinct CRC organoid (PDOs) models (TX-CC-201, TX-CC-199, TX-CC-286, TX-CC-096h2, TX-CC-208, TX-CC-206) to evaluate drug response in this study. Cells were seeded at a 1×10^4^ cells/well in 6-well plates and exposed to a serial drug concentration range (62.5 nM – 4000 nM) as indicated in **Fig. 1**, and total organoid number and growth were assessed after 10 to 12 days of treatment. Representative brightfield images demonstrate a progressive, dose-dependent suppression of organoid growth (size, number, or area). Notably, CF10 treatments exhibited visibly stronger growth inhibition compared to matched 5-FU dosages across six independent models (**Fig. 1A-L**).

**Figure 1.**
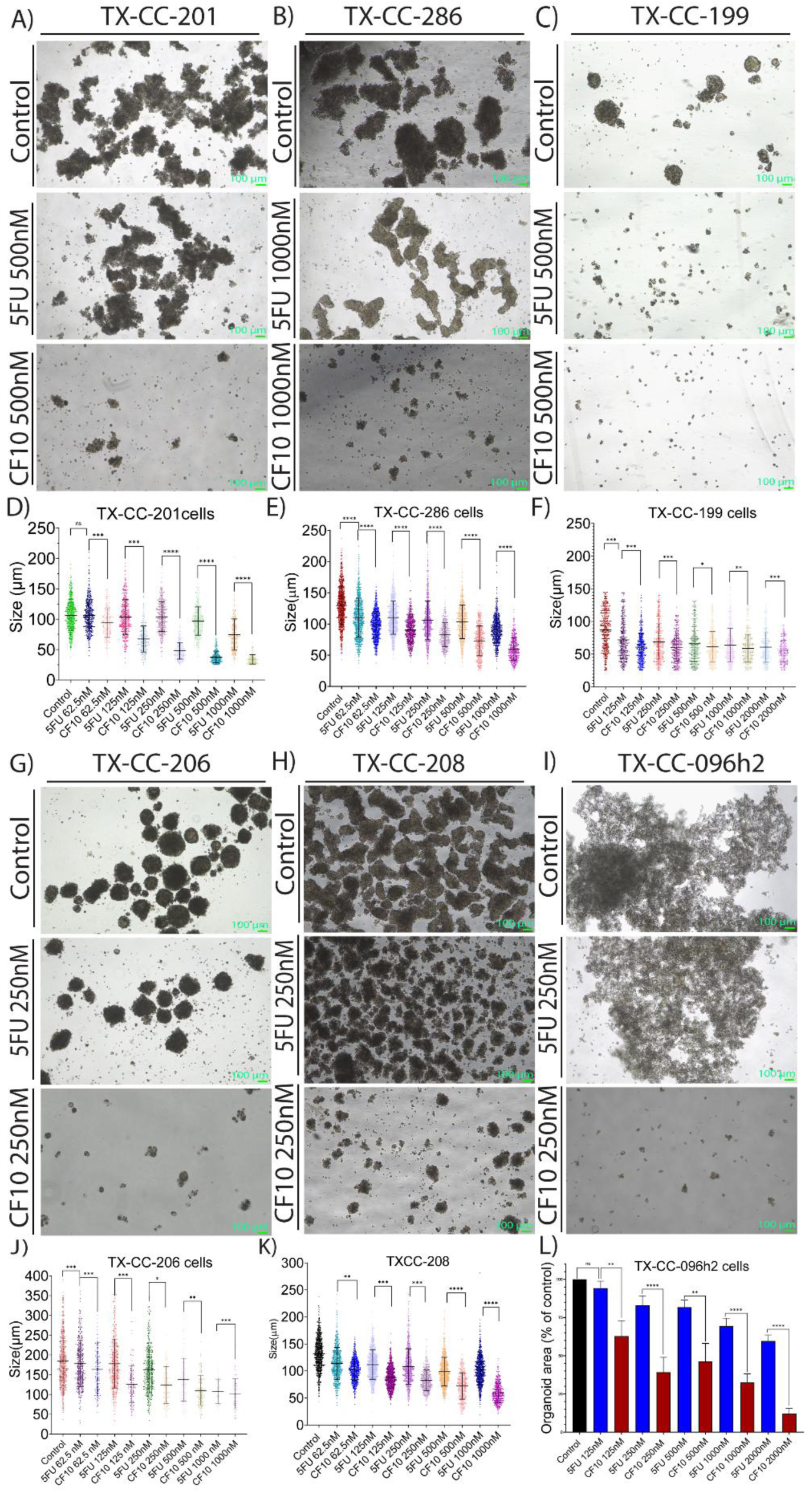
CF10 suppresses long-term growth of patient-derived CRC organoids across diverse models under normoxia. Patient-derived colorectal cancer organoids (PDOs) representing distinct morphologies were treated for 10–12 days with vehicle (Control), 5-fluorouracil (5-FU), or CF10 at the indicated concentrations. **(A–C, G–I)** Representative phase-contrast images of TX-CC-201, TX-CC-286, TX-CC-199, TX-CC-206, TX-CC-208, and TX-CC-096h2 organoids following treatment (dose shown on left). **(D–F, J–K)** Quantification of organoid size distributions for the indicated PDO models across the dose ranges shown on the x-axes; each point represents an individual organoid, and bars indicate mean ± SD. **(L)** Quantification of total organoid area per imaging field for TX-CC-096h2 (sheet/tubule morphology), normalized to control, across equimolar 5-FU and CF10 concentrations. Statistical significance is indicated (*P<0.05, **P<0.01, ***P<0.001, \*\*\**P<0.0001*); replicate structure, and statistical tests are described in Methods.

Quantification of organoid size (μm) demonstrated that CF10 caused significantly greater reductions in size than 5-FU at matched molar concentrations (**Fig. 1A-C, G–I**; P<0.05, unpaired t-tests). In the TX-CC-096h2 model, which uniquely forms spreading sheet- and tubule-like structures, growth was quantified as total organoid area per field (μm^2^). Even in this atypical growth architecture, CF10 produced a more pronounced, dose-dependent reduction in area compared to 5-FU (Welch t-test). Collectively, these findings demonstrate that CF10 maintains potent anti-tumor activity compared to 5-FU regardless of morphological heterogeneity, suggesting its potential to overcome the physiological barriers that frequently limit the efficacy of conventional fluoropyrimidines.

### Physiological oxygen or physioxia (5% O_2_) reveals morphological plasticity of PDOs and confirms CF10 efficacy

Standard ∼20% O_2_ culture conditions are significantly hyperoxic compared to the ambient low oxygen tensions characteristic of the solid tumor microenvironment. This discrepancy is critical, as oxygen availability is known to modulate cellular signaling, metabolic phenotypes, and the resulting sensitivity to chemotherapeutics.^39,40^ Considering these variables, we evaluated organoid responses under physioxia (5% O_2_) to determine if CF10’s therapeutic advantage over 5-FU persists in a more clinically/physiologically representative context. We used three different PDO models (TX-CC-096h2, TX-CC-286, and TX-CC-199) that were able to grow in these conditions. We observed significant phenotypic plasticity; specifically, the TX-CC-286 model exhibited a striking oxygen-dependent morphological shift, transitioning from the compact, rounded spheroids (**Fig. 1B**) seen in normoxia to expansive, tubule-like architectures (**Fig. 2B**). Despite these profound organoid structural differences, CF10 maintained significantly more potent growth suppression than equimolar 5-FU across the entire concentration range (**Fig. 2D-F**). Quantification of organoid size/area and number confirmed that the superior efficacy of CF10 was not attenuated by the low-oxygen environment. Collectively, these findings suggest that the therapeutic advantage of CF10 is robust across clinically relevant oxygen contexts and remains uncompromised by physioxia-induced phenotypic remodeling, reinforcing its potential efficacy within the poorly oxygenated regions of solid CRC tumors.

**Figure 2.**
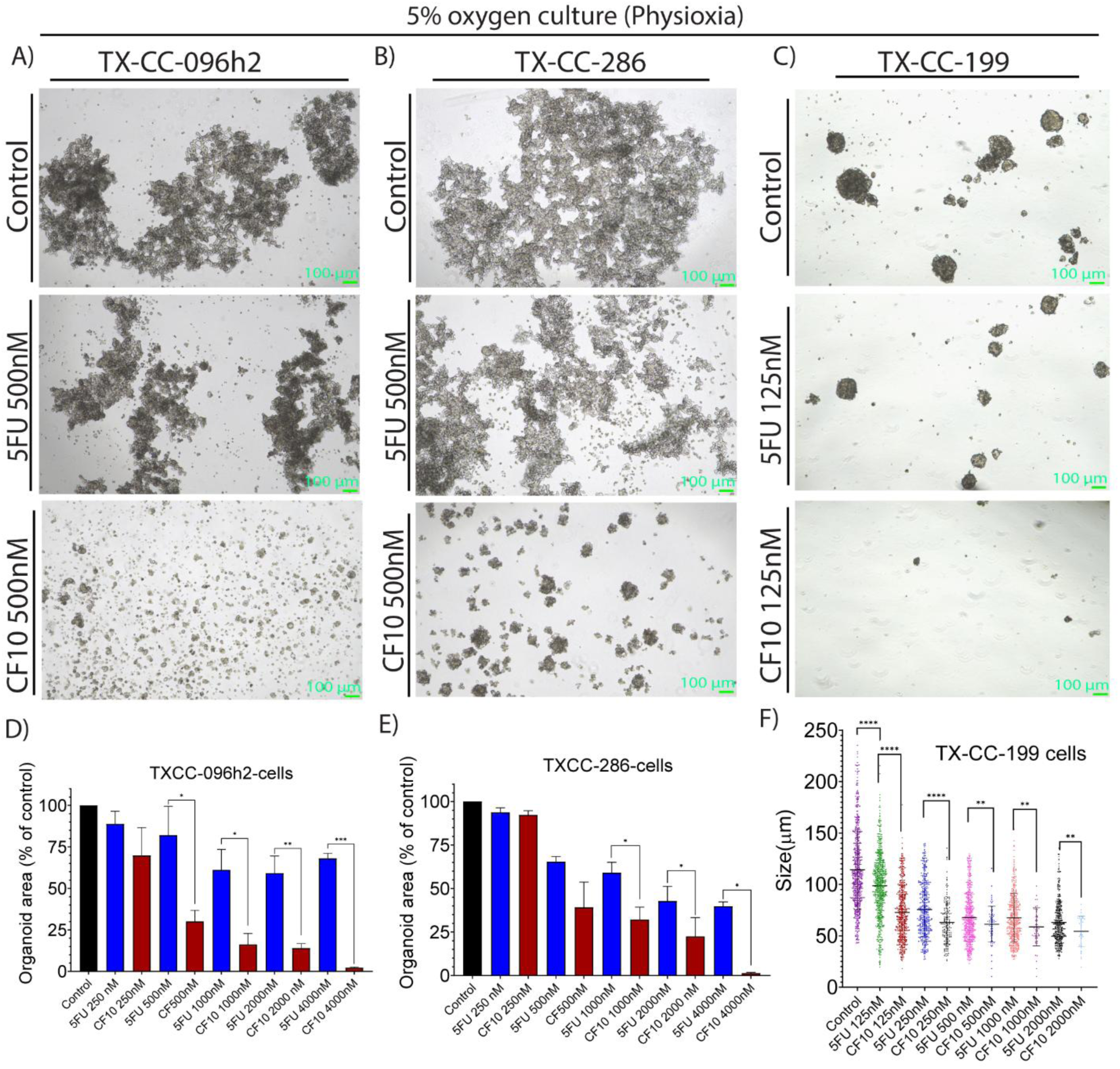
CF10 retains superior anti-organoid activity under physioxia (5% O₂). Patient-derived colorectal cancer organoids (TX-CC-096h2, TX-CC-286, and TX-CC-199) were cultured under physioxia (5% O₂) and treated for 10–12 days with vehicle (Control), 5-fluorouracil (5-FU), or CF10 at the indicated concentrations. **(A–C)** Representative phase-contrast images of TX-CC-096h2, TX-CC-286, and TX-CC-199 organoids following treatment. **(D–E)** Total organoid area per imaging field (normalized to control) for TX-CC-096h2 and TX-CC-286 across equimolar 5-FU and CF10 doses. Bars represent mean ± SEM. **(F)** Organoid size distribution for TX-CC-199 across the indicated dose conditions under physioxia; each point represents an individual organoid, and horizontal bars indicate mean ± SD. Statistical comparisons between 5-FU and CF10 at each dose were performed using Welch’s t-test; significance is indicated as *P<0.05, **P<0.01, ***P<0.001, \*\*\**P<0.0001*.

### CF10 induces stronger DNA damage/replication stress signaling and cell death in comparison to 5-FU in CRC PDOs

Efficacy of fluoropyrimidine therapy depends on the induction of lethal DNA damage and the activation of programmed cell death. Therefore, we assessed pharmacodynamic responses by measuring γH2AX (H2AX Ser139 phosphorylation) and Top1cc (topoisomerase I–DNA cleavage complexes) as markers of DNA damage/Top1 poisoning, Ki67 as a proliferation marker, and Annexin V/PI staining to quantify apoptotic and necrotic cell populations.^41,42^ To examine the early molecular consequences of treatment, TX-CC-199 and TX-CC-201 organoid cultures were exposed to equimolar concentrations (1 μM) of CF10 and 5-FU. Immunoblotting revealed that both treatments induced γH2AX phosphorylation compared to control cultures, and specifically, CF10 elicited a markedly stronger signal than 5-FU in both organoid models, indicating a more robust DNA damage stress induction by CF10 (**Fig. 3G, H**). Similarly, monoubiquitination of FANCD2 (**Fig. 3G, H**) indicates repair of DNA damage, specifically highlighting the presence of stalled replication forks and, crucially, DNA double-strand breaks (DSBs). Consistent with the previous studies^15^, these damages were also accompanied by significantly higher Top1cc formation (**Fig. 3B**) and pRPA32-S33 (**Fig. 3C**) immunofluorescence (IF) intensities. These findings were further validated via significant elevation of IF intensities for γH2AX (Fig. 3A) and the proliferation marker Ki-67 (Fig. 3I). Consistent with the immunoblot data, IF quantification demonstrated significantly higher levels of γH2AX foci and a concomitant reduction in Ki-67 following CF10 treatment compared to 5-FU.

**Figure 3.**
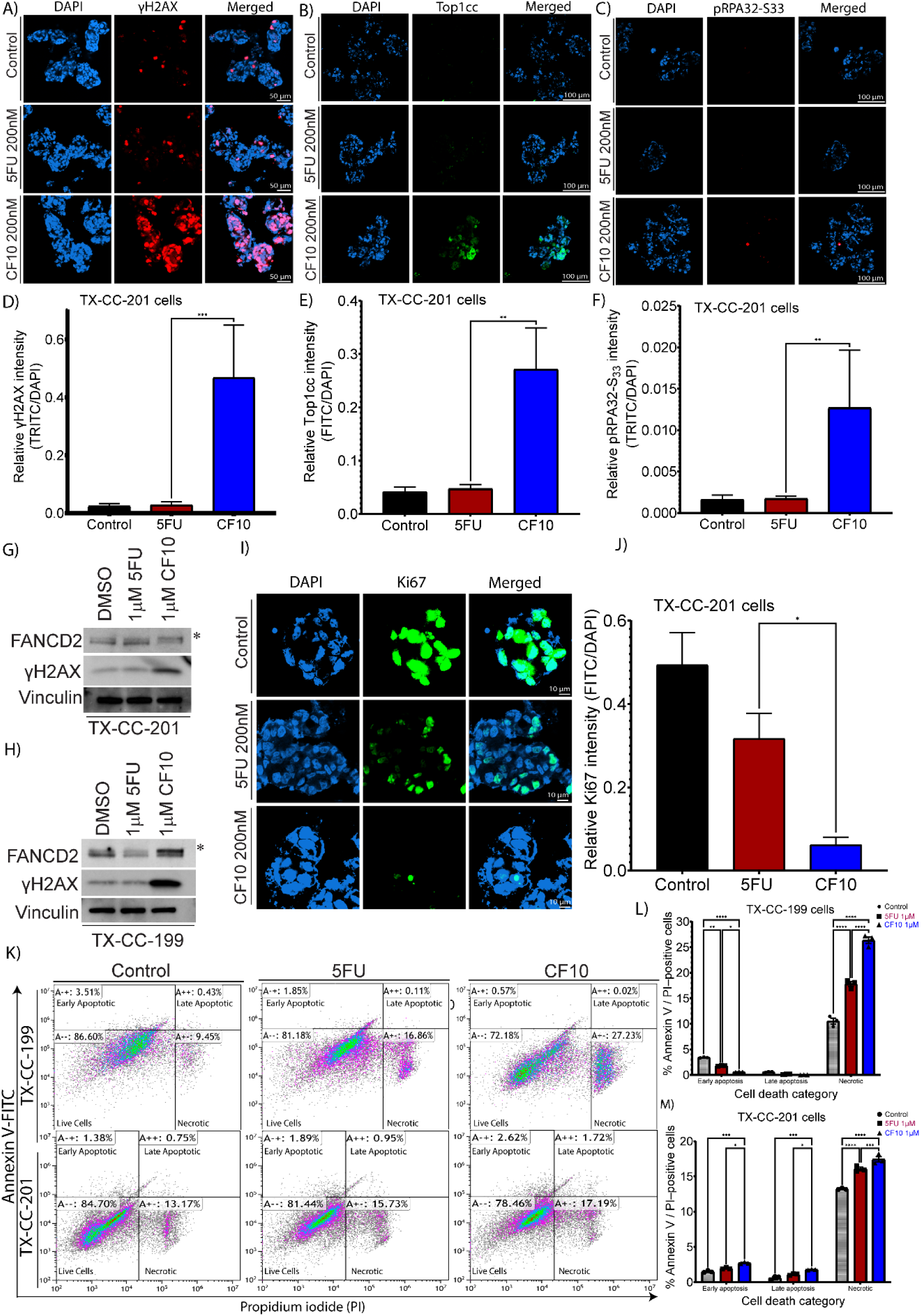
CF10 induces stronger DNA damage/replication-stress signaling, reduced proliferation, and increased terminal cell death than 5-FU in CRC PDOs. TX-CC-199 and TX-CC-201 patient-derived CRC organoids were treated with vehicle (Control), 5-fluorouracil (5-FU), or CF10 at the indicated concentrations. **(A–C)** Representative immunofluorescence images from TX-CC-201 organoids showing DAPI and staining for γH2AX (Ser139) (A), Top1cc (B), and pRPA32-S33 (C), with merged channels (48 hrs). **(D–F)** Quantification of γH2AX, Top1cc, and pRPA32-S33 fluorescence intensity normalized to DAPI in TX-CC-201 organoids. **(G–H)** Immunoblot analysis of FANCD2 and pH2AX-S139 in TX-CC-201 (G) and TX-CC-199 (H) organoids treated with equimolar 5-FU or CF10 (1 μM, 24 hrs); vinculin served as a loading control. **(I–J)** Representative Ki67 immunofluorescence images (I) and quantification of Ki67 intensity normalized to DAPI (J) in TX-CC-201 organoids (48 hrs). **(K)** Representative Annexin V/PI flow cytometry plots from TX-CC-199 and TX-CC-201 organoids following 72 hrs treatment. **(L–M)** Quantification of Annexin V/PI-defined populations (early apoptosis, late apoptosis, and PI-positive/necrotic) for TX-CC-199 (L) and TX-CC-201 (M). For panels **D–F** and **J**, statistical comparisons were performed using two-tailed Mann–Whitney tests. For panels **L–M**, differences among treatments were assessed by two-way ANOVA with Tukey’s multiple comparisons. Significance is indicated as *P<0.05, **P<0.01, ***P<0.001, \*\*\**P<0.0001*.

We next assessed the treatment-associated cell death phenotypes in these models using Annexin V/PI staining followed by flow cytometry at 72 hours of drug exposure. In TX-CC-199 organoids, both 5-FU and CF10 treatments exhibited increased cell death phenotype relative to controls (**Fig. 3K-M**). Two-way ANOVA with Tukey’s multiple comparisons confirmed that CF10 significantly increased early apoptosis compared to both 5-FU (P<0.0001) and control (P<0.0001). Notably, CF10 produced a significantly greater PI-positive fraction than 5-FU (P=0.0008), suggesting stronger terminal cytotoxicity. Interestingly, the lower portion of early apoptotic cells observed with CF10 in the TX-CC-199 model, despite the marked increase in PI-positivity, likely indicates a rapid progression from early apoptosis into late-stage secondary necrosis by the 72-hour timepoint. Whereas in TX-CC-201, drug treatments induced similar shifts, with CF10 again outperforming 5-FU most prominently within the PI-positive compartment **(Fig. 3K**). CF10 significantly increased early apoptosis (vs. 5-FU, P=0.0196) and late apoptosis (vs. 5-FU, P=0.0339), while markedly expanding the PI-positive fraction compared to both control and 5-FU (both P<0.0001). Representative flow cytometry plots illustrate these distinct distributional changes across both models (**Fig. 3L, M**). Collectively, these data demonstrate that CF10 elicits a more potent cytotoxic response than equimolar 5-FU in patient-derived CRC models. While cell-line-specific differences exist in the temporal distribution of death states, the overarching trend confirms that CF10 more effectively triggers DNA damage and drives cells toward terminal death.

### CF10 potently suppresses the invasive outgrowth of CRC PDOs even at 10-fold lower doses than 5-FU

Local invasion is a hallmark capability that underlies metastatic progression and poor outcomes in colorectal cancer. Consequently, 3D invasion assays provide a clinically relevant functional endpoint that extends beyond simple growth inhibition.^43,44^ Because these invasive states are frequently linked to stem-like properties^45,46^, we investigated whether CF10’s superior cytotoxicity extends to these high-risk functional phenotypes. To determine whether CF10 more effectively restricts invasive outgrowth, we quantified the invasion area of TX-CC-096h2 organoids over seven days using two dosing schemes (**Fig. 4A-F**): an equimolar comparison (5 nM) and a potency-shifted comparison (3 nM CF10 vs. 30 nM 5-FU). Importantly, concentrations were deliberately chosen within a low-dose range, including minimally growth-inhibitory conditions, to minimize the possibility that reduced invasive outgrowth simply reflected generalized cytotoxicity. While invasion areas were uniform across all groups at baseline (Day 0), both fluoropyrimidines significantly suppressed (**Fig. 4E, F**) invasive expansion relative to control by Day 3, with CF10 eliciting the most robust inhibition. In the equimolar comparison (5 nM), CF10-treated organoids exhibited significantly less invasion than those treated with 5-FU at both Day 3 (5-FU–CF10: 95% CI 12.63–25.37, P<0.0001) and Day 7 (P<0.0001; **Fig. 4A**).

**Figure 4.**
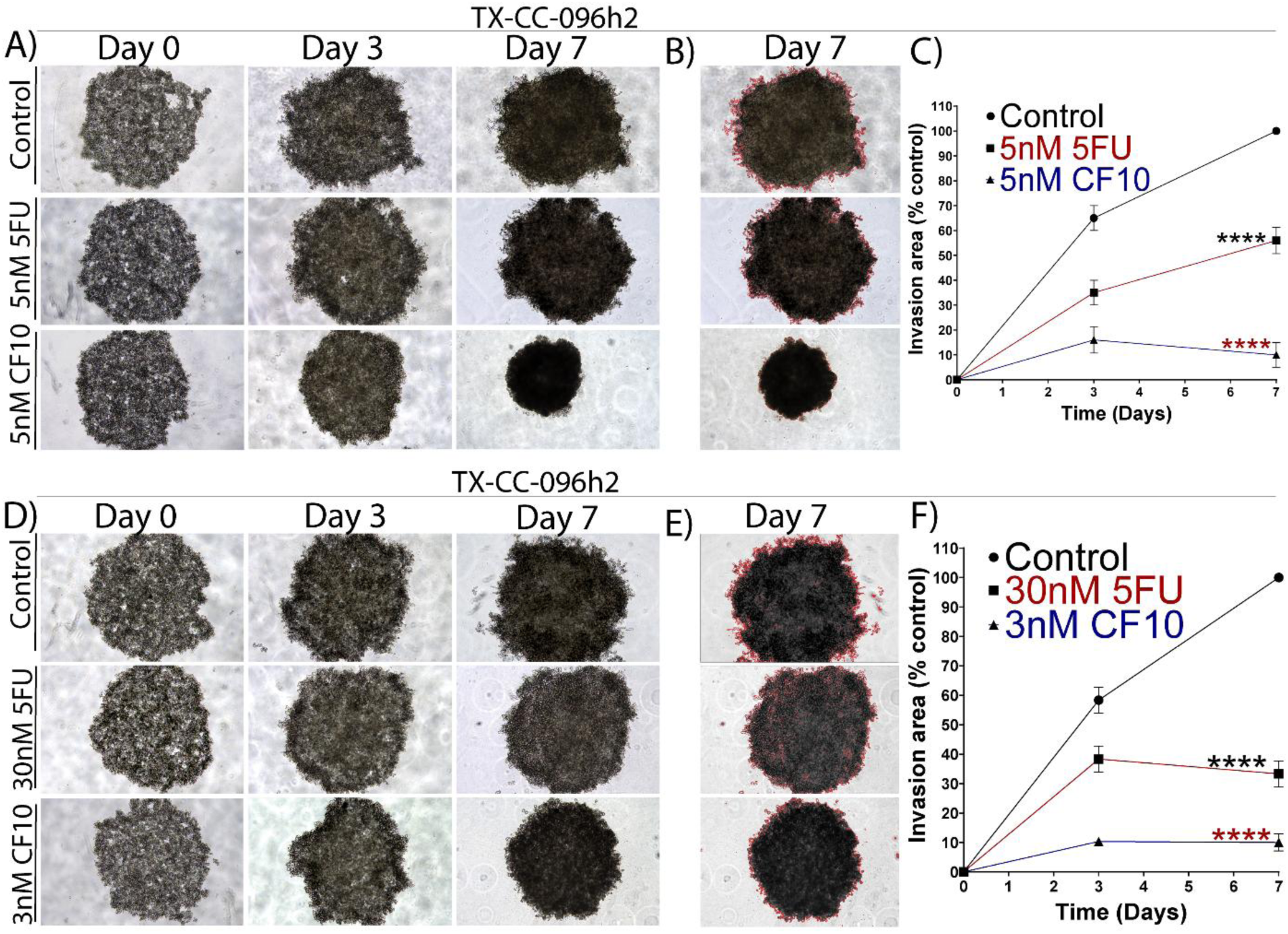
CF10 suppresses 3D invasion of TX-CC-096h2 CRC organoids more effectively than 5-FU, including at a one-log lower molar dose. TX-CC-096h2 patient-derived CRC organoids were embedded in 3D Matrigel and monitored for invasive outgrowth over time under vehicle (Control), 5-FU, or CF10 treatment. **(A)** Representative phase-contrast images at days 0, 3, and 7 comparing equimolar dosing (5 nM 5-FU vs 5 nM CF10). **(B)** Representative images at days 0, 3, and 7 comparing one-log lower CF10 dosing (30 nM 5-FU vs 3 nM CF10). **(C–D)** Day 7 invasion outlines (red) used to quantify the invasive area for the corresponding dosing conditions in (A) and (B). **(E–F)** Quantification of invasion area over time (normalized to control) for equimolar dosing (E) and one-log lower CF10 dosing (F). Data are shown as mean ± SEM (n = 3 independent replicates per condition). Statistical comparisons at each time point were performed using two-way ANOVA with Šidák’s multiple comparisons (within each time point); significance is indicated as *P<0.05, **P<0.01, ***P<0.001, \*\*\**P<0.0001*.

Remarkably, CF10 maintained superior anti-invasive activity even in the one-log potency-shifted condition (**Fig. 4B, D, F**). Despite being administered at a 10-fold lower concentration, 3 nM CF10 significantly outperformed 30 nM 5-FU at both Day 3 (30 nM 5-FU–3 nM CF10: 95% CI 17.89–38.11, P<0.0001) and Day 7 (95% CI 13.22–33.45, P<0.0001; **Fig. 4F**). Together, these data demonstrate that CF10 more potently suppresses 3D invasive expansion than 5-FU, including under low-dose conditions designed to minimize confounding by generalized cytotoxicity, thereby supporting a true anti-invasive effect beyond simple dose-dependent growth suppression, thus, providing enhanced functional control of invasion-linked phenotypes in patient-derived models.

### CF10 strongly depletes ALDH-positive stem-like cells compared with 5-FU

Invasive outgrowth in 3D culture often reflects the acquisition of motile programs associated with epithelial–mesenchymal plasticity.^45,47,48^ Given that these states are frequently linked to stem-like properties^46,47^, we next investigated whether these treatments modulate the cancer stem cell (CSC) compartment. We quantified aldehyde dehydrogenase high (ALDH-high) population using the ALDEFLUOR assay, utilizing DEAB-treated controls to strictly define the ALDH-low gates. In both models, untreated controls maintained a substantial ALDH-positive fraction (TX-CC-096h2: ∼22%; TX-CC-199: ∼19%). In TX-CC-096h2, 5-FU failed to significantly reduce the ALDH-positive population relative to control (P=0.2935). In contrast, CF10 markedly decreased ALDH positivity compared to both control (P<0.0001) and 5-FU (P=0.0001; **Fig. 5A**). A similar trend was observed in TX-CC-199; while both drugs reduced ALDH positivity, CF10 again elicited the most robust suppression (5-FU vs. control, P=0.0038; CF10 vs. control, P=0.0001). Crucially, CF10 was significantly more effective than 5-FU at depleting this population (P=0.0479; **Fig. 5A-C**). Collectively, these data indicate that CF10 more effectively depletes the ALDH-high, stem-like subpopulation - a compartment frequently linked to chemoresistance and disease relapse in CRC.^49^ This depletion complements the superior anti-invasive activity observed in our 3D assays and suggests that CF10 may overcome 5-FU resistance by targeting the CSC niche.

**Figure 5.**
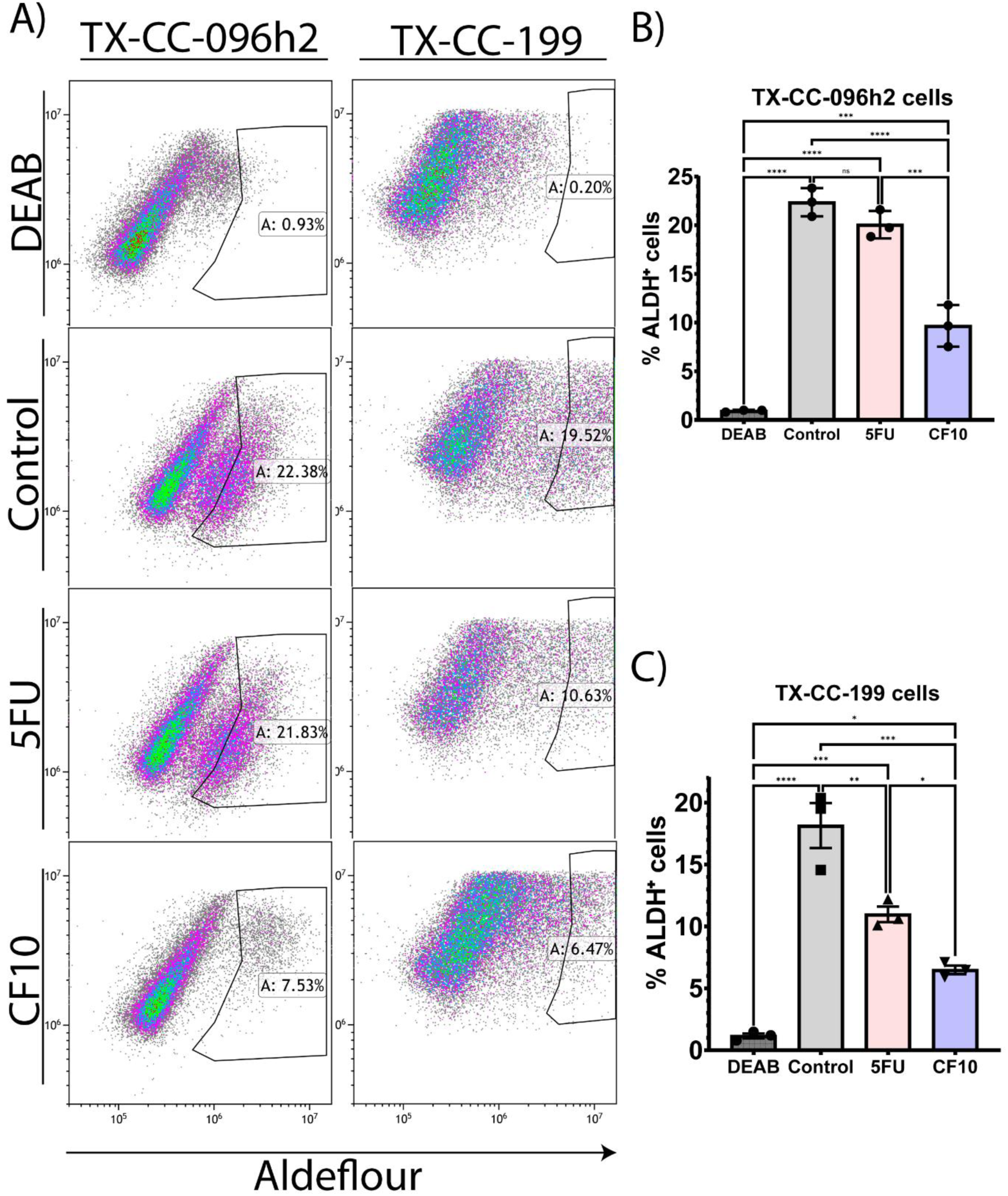
CF10 reduces the ALDH-high stem-like compartment compared with 5-FU in patient-derived CRC models. PDO-derived single cells from TX-CC-096h2 and TX-CC-199 were treated with vehicle (Control), 5-fluorouracil (5-FU), or CF10 (1 μM, 72 hrs) and ALDH activity was quantified by ALDEFLUOR™ flow cytometry. **(A)** Representative ALDEFLUOR™ flow cytometry plots and gating strategy for each model; DEAB-treated samples (ALDH inhibitor) were used to define the ALDH^high gate. **(B–C)** Quantification of the ALDH^high fraction (% ALDH^+ cells) in TX-CC-096h2 (B) and TX-CC-199 (C). Bars represent mean ± SEM with individual points indicating biological replicates (n = 3 per condition). Statistical significance was assessed by ordinary one-way ANOVA with Tukey’s multiple comparisons; *P<0.05, **P<0.01, ***P<0.001, \*\*\**P<0.0001*.

### In vivo efficacy of CF10 in patient-cell-derived xenograft (PCDX) models

To validate our *in vitro* findings in a more complex physiological environment, we transitioned CF10 into PCDX models that better recapitulate the architecture of human CRC. 5-FU-based regimens remain a core component of metastatic colorectal cancer therapy; however, incomplete responses and resistance are common. This creates a need for improved fluoropyrimidine strategies that retain activity in clinically relevant, patient-derived models.^35,50,51^ We, therefore, evaluated the efficacy of CF10 versus 5-FU using PCDX models generated from patient-derived primary cells. The TX-CC-199 and TX-CC-201 lines were selected for *in vivo* validation as they represent distinct patient-derived backgrounds that demonstrated differential apoptotic and stem-like profiles in our 3D *in vitro* assays. Following PDX engraftment and processing into single-cell suspensions, PCDX tumors were established in immunodeficient mice (**Fig. 6A**).

**Figure 6.**
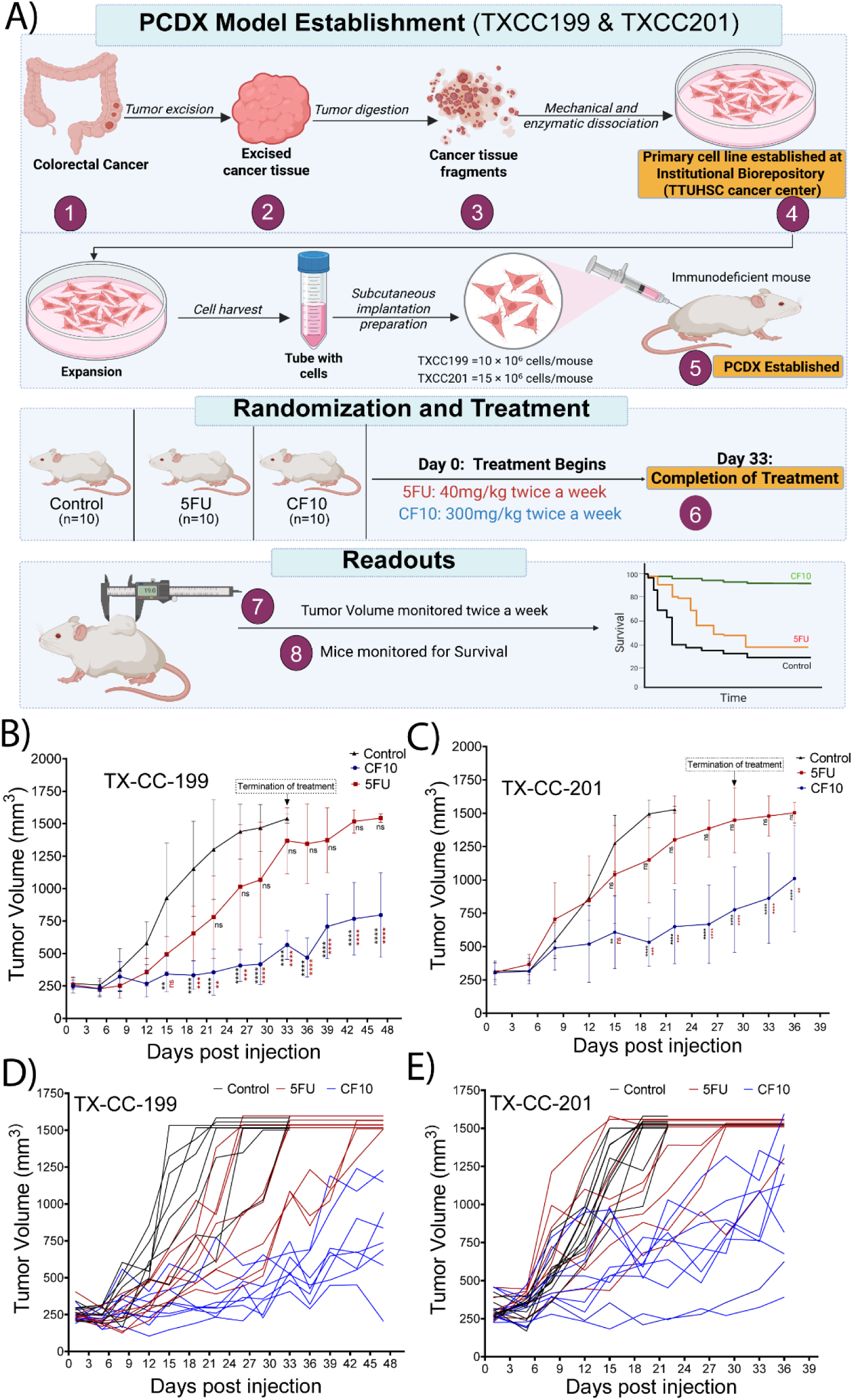
CF10 improves tumor control and survival in patient cell–derived xenograft models of CRC. **(A)** Schematic of model establishment and in vivo study design for TX-CC-199 and TX-CC-201 patient cell–derived xenografts (PCDX). Tumor tissue was processed to generate a primary cancer cell suspension, expanded, and implanted subcutaneously into immunodeficient mice to establish CDX models. Mice were randomized into treatment groups (Control, 5-FU, CF10; n = 10 per group). Treatment began on day 0 and continued through day 33: 5-FU (40 mg/kg, twice weekly) or CF10 (300 mg/kg, twice weekly). Tumor volume was measured twice weekly, and mice were monitored for survival. **(B–C)** Mean tumor volume over time (mm³; mean ± SEM) in TX-CC-199, n=8 mice per arm, 2 mice/treatment group were excluded as outliers (B) and TX-CC-201, n=9 mice per arm, 1 mouse/treatment group was excluded as an outlier (C) xenografts for Control, 5-FU, and CF10; the treatment period is indicated. **(D–E)** Individual tumor growth trajectories for TX-CC-199 (D) and TX-CC-201 (E) showing mouse-level responses within each treatment group. Statistics: Tumor volumes were compared between groups at each time point using multiple unpaired t tests (one test per time point) with Holm–Šidák correction for multiple comparisons (α = 0.05), including CF10 vs Control, 5-FU vs Control, and CF10 vs 5-FU.

Mice were implanted subcutaneously with TX-CC-199 (10×10^6^ cells) or TX-CC-201 (15×10^6^ cells) and randomized (n=10 per arm) to receive vehicle control, 5-FU (40 mg/kg, twice weekly), or CF10 (300 mg/kg, twice weekly) from Day 0 through Day 33 (**Fig. 6A**). Across both models, CF10 produced more robust suppression of tumor growth than 5-FU or control (**Fig. 6B–C**). The divergence between CF10 and 5-FU became increasingly evident as tumors progressed toward the end of the dosing window, with significant statistical separation at terminal timepoints (Day 33). Individual mouse trajectories further corroborated these group-level effects: CF10 consistently shifted tumor growth curves downward and reduced the frequency of rapid outgrowth compared with both 5-FU and control (**Fig. 6D–E**). Collectively, these data demonstrate that CF10 maintains superior antitumor activity in heterogeneous primary CRC-derived *in vivo* models.

### CF10 improves survival in patient-derived CRC models without overt toxicity

The superior tumor control observed with CF10 translated into a significant survival advantage across multiple patient-derived backgrounds. In both the TX-CC-199 and TX-CC-201 models, we were able to evaluate survival across models with differing growth kinetics. Kaplan–Meier analysis demonstrated that CF10 produced a clear survival benefit in both cohorts (**Fig. 7A, C**; **Table 1**).

**Figure 7.**
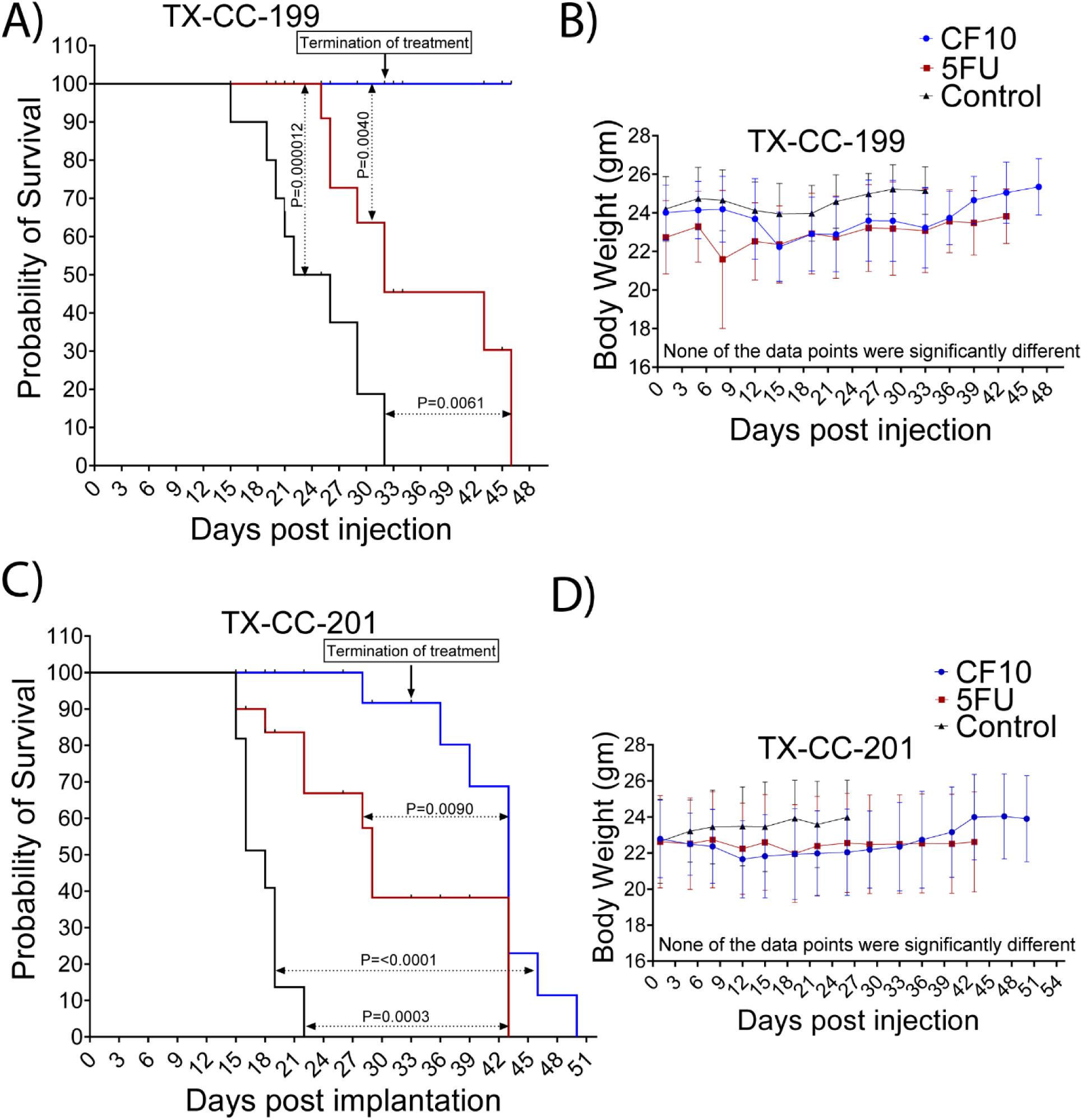
CF10 improves survival without significant body weight loss in CRC patient cell–derived xenograft models. **(A, C)** Kaplan–Meier survival curves for mice bearing TX-CC-199 (A) or TX-CC-201 (C) xenografts treated with vehicle (Control), 5-fluorouracil (5-FU), or CF10 according to the regimen in Fig. 6; treatment termination is indicated. Survival differences were assessed by the log-rank (Mantel–Cox) test, with P values shown. **(B, D)** Body weight trajectories (mean ± SEM) for TX-CC-199 (B) and TX-CC-201 (D) cohorts during treatment and follow-up. Body weight was analyzed using a mixed-effects model (REML) with time and treatment as fixed effects; no significant treatment effect was detected in either cohort (TX-CC-199: P = 0.0888; TX-CC-201: P = 0.5088).

**Table 1.** Summary of median survival for each model and treatment arm, with log-rank P values for CF10 vs 5-FU and cohort sizes (n/arm).

| CDX model | Median survival (days) |  |  | Log-rank P<br>(CF-10 vs 5-FU) | n/arm |
| --- | --- | --- | --- | --- | --- |
|  | Control | 5FU | CF10 |  |  |
| TXCC201 | 18 | 29 | 43 | 0.00990 | 9 |
| TXCC199 | 24 | 32 | NR | 0.0040 | 8 |

In TX-CC-201, CF10 extended median survival to 43 days compared with 29 days for 5-FU and 18 days for control (log-rank P=0.0099). In TX-CC-199, CF10 yielded an even more durable benefit; while median survival was 32 days for 5-FU and 24 days for control, the CF10 group did not reach (NR) the median survival during the 50-day observation window (P=0.0040 vs. 5-FU). To assess tolerability, we tracked longitudinal body weight throughout treatment and follow-up (**Fig. 7B, D**). Body weights remained comparable across all groups, with no evidence of treatment-related weight loss or physical distress in the CF10 arms. Together, the significant survival gains and stable body weights support the conclusion that CF10 improves *in vivo* outcomes in patient-derived CRC models without compromising systemic tolerability under the tested dosing schedule.

## Discussion

In this study, we utilized primary patient-derived CRC models, including 3D organoid cultures, to evaluate the efficacy of the next-generation fluoropyrimidine, CF10. By leveraging organoids that recapitulate the architecture and heterogeneity of patient tumors^52^, we established a clinically relevant platform that captures tumor-specific resistance features more accurately than traditional 2D systems. This approach underscores the translational importance of our findings, demonstrating that CF10 maintains markedly superior anti-tumor activity compared to 5-FU, even in models that are 5-FU-refractory.

Across our six PDO models, we observed their limited responsiveness towards 5-FU across the tested dose range, indicating a functionally 5-FU-recalcitrant phenotype in this 3D setting. In contrast, CF10 produced consistent and substantially greater inhibition of organoid growth across the same panel, demonstrating that its activity is maintained even when conventional 5-FU shows little effect. This pattern is clinically relevant because fluoropyrimidine response in CRC is highly heterogeneous and is shaped by both intrinsic tumor biology and treatment history, including selection for resistant subclones during prior fluoropyrimidine exposure^53,54^ Importantly, PDOs are increasingly used as “living biomarkers” that preserve patient-specific, tumor-intrinsic determinants of drug response, and multiple studies support their utility for modeling chemotherapy sensitivity/resistance in CRC and related settings.^55–57^ A key limitation of our current dataset is that we lack detailed, standardized clinical annotation (e.g., prior 5-FU exposure, metastatic vs primary origin, interval from therapy, and molecular features such as TS/DPD (dihydropyrimidine dehydrogenase)-related fluoropyrimidine metabolism and DDR context) that could explain why these PDOs were broadly 5-FU–unresponsive. Future studies integrating comprehensive patient treatment histories and molecular stratification with PDO pharmacotyping will be important to distinguish intrinsic versus acquired fluoropyrimidine resistance and to identify predictive correlates of CF10 sensitivity and determine whether CF10 bypasses or directly overcomes the same biologic programs that blunt 5-FU response.

Efforts to overcome 5-FU resistance have historically focused on regimen intensification, such as the addition of oxaliplatin or irinotecan.^50,58,59^ While these combinations can extend disease control, they are often limited by a narrow therapeutic window and cumulative toxicities.^60,61^ Another central translational limitation of conventional 5-FU is that effective tumor control depends on generating sufficiently strong and sustained intracellular stress, yet clinical dosing is constrained by toxicity and pharmacologic inefficiency. In that context, CF10 can be viewed not simply as a more potent fluoropyrimidine but as a fluoropyrimidine redesign intended to improve the quality and persistence of tumor-directed stress without relying on greater systemic exposure.^62,63^ Mechanistically, CF10 is engineered to bypass these constraints via a dual mechanism of action: it directly releases the active metabolite FdUMP to inhibit TS while concurrently stabilizing Top1-DNA cleavage complexes.^54^ This multifaceted design allows CF10 to retain potent cytotoxicity in contexts where 5-FU fails due to elevated TS expression or impaired metabolic activation.^64^

Our pharmacodynamic profiling in patient-derived CRC organoids supports a model in which CF10 triggers a quantitatively stronger genotoxic stress than equimolar 5-FU. Across two independent PDO models, CF10 produced significantly more robust γH2AX than 5-FU, which is consistent with activation of intensified DNA damage response and associated chromatin signaling and broader DDR response, suggesting recruitment and amplification of repair pathways at sites of DNA damage.^41,65^ Importantly, this elevated DDR response was accompanied by increased Top1cc signal and elevated pRPA32-S33, linking CF10 exposure to formation of topoisomerase I–DNA cleavage complexes and activation of ATR-associated replication stress signaling. RPA32 Ser33 phosphorylation is a well-established marker of ATR pathway activation in response to Top1 poison–associated lesions and replication-associated DNA damage, providing a mechanistic bridge between Top1-linked damage and downstream checkpoint signaling.^66^ Moreover, FANCD2 is a central effector of the Fanconi anemia pathway that is recruited to stalled/collapsed replication forks and coordinates fork protection and lesion resolution during replication stress.^67^ Consistent with this biology, increased FANCD2 in CF10-treated PDOs supports the interpretation that CF10 drives replication-coupled lesions that engage FA/ATR checkpoint pathways beyond what is observed with equimolar 5-FU.^66^ Together, these convergent readouts argue that CF10 produces a more intense lesion burden and/or more persistent damage processing than 5-FU in a 3D patient-derived context.

Because local invasion underlies progression, dissemination, and eventual loss of curative treatment opportunities in CRC, the greater suppression of invasive outgrowth by CF10 may have translational relevance beyond bulk tumor control, although our assay does not directly model resectability.^68,69^ Our functional assays revealed that CF10 effectively suppresses 3D invasive outgrowth and depletes the ALDH-high stem-like population. Invasive capacity and stemness often co-evolve through epithelial–mesenchymal plasticity (EMP) programs.^70,71^ While we do not infer a direct causal link between these two phenotypes, the convergent reduction in both invasion and ALDH activity indicates that CF10 targets the aggressive, treatment-refractory cell states that drive clinical relapse.^31,72^ Since ALDH-high populations are enriched for tumor-initiating cells and associated with multidrug resistance^30,49^, the ability of CF10 to more effectively clear this niche compared to 5-FU suggests a potential to reduce the risk of disease recurrence.

A critical translational component of this work was the *in vivo* validation using PCDX models. While PDX models offer high complexity, PCDX models provide the reproducibility and standardized growth kinetics necessary for rigorous, head-to-head efficacy and survival comparisons.^73^ By using these controlled platforms, we demonstrated that CF10’s efficacy advantage translates to superior tumor control and a robust survival benefit without an associated “tolerability penalty.” The stability of body weights across treatment groups suggests that CF10 achieves its enhanced on-target effects without the metabolic liabilities or catabolite-driven toxicities typically associated with 5-FU.^15^

One point that merits clarification is the difference between the administered doses of 5-FU and CF10. These regimens are not directly comparable on a simple mg/kg basis, because CF10 is not a free 5-FU but a polymeric fluoropyrimidine whose additional structural components influence molecular mass, pharmacology and tolerability. Importantly, the 5-FU schedule used here (40 mg/kg, twice weekly) falls within the range of active regimens reported in preclinical CRC models, supporting the view that our comparator arm was pharmacologically relevant rather than underdosed. This dosing strategy was also informed by our prior CF10 studies. In an orthotopic CRC model, higher-intensity 5-FU treatment was associated with intestinal crypt apoptosis, neutrophilic infiltration, and edema, whereas CF10 showed normal blood chemistry parameters and reduced gastrointestinal (GI) tissue injury despite superior antitumor activity.^15^ In a separate orthotopic rat colorectal liver metastasis study, CF10 likewise showed superior tolerability and antitumor efficacy relative to 5-FU, without treatment-associated weight loss^13^ Taken together with the present findings, namely significantly greater tumor burden reduction, significantly prolonged survival, and no significant body-weight penalty, these data support the conclusion that CF10 provides a broader therapeutic window/index relative to 5-FU, whose efficacy is more tightly constrained by the narrower tolerability limits of conventional fluoropyrimidine therapy.

Importantly, these findings should be interpreted as a structured escalation of translational evidence rather than a series of isolated assay-level observations, because CF10 consistently remained favorable across heterogeneous PDOs, physioxic culture, invasion/stem-like endpoints, and *in vivo* validation. Collectively, these findings provide a compelling rationale for the clinical development of CF10. By maintaining high efficacy in 5-FU-resistant primary CRC models and demonstrating a favorable safety profile *in vivo*, CF10 addresses a major unmet need for patients who have progressed on standard-of-care regimens like FOLFOX or FOLFIRI, ultimately becoming recalcitrant. This work establishes a robust translational framework, integrating patient-derived organoids with reproducible *in vivo* assays, to support the advancement of CF10 into Phase I clinical trials.

## Conclusion

This study provides a rigorous head-to-head evaluation of CF10 versus 5-FU across multiple distinct patient-derived CRC models. By linking molecular DNA damage signaling to 3D functional suppression and validating these results with survival endpoints in two independent PCDX cohorts, we position CF10 as a promising next-generation fluoropyrimidine candidate. As 5-FU remains the cornerstone of CRC therapy, the translational development of agents like CF10 that can overcome acquired resistance in clinically relevant models is essential for improving long-term patient outcomes.

## Supporting information

Supplementary Macro S1

## Acknowledgments

We thank the TTUHSC Cancer Center and C. Patrick Reynolds, M.D., Ph.D., for providing the patient-derived colorectal cancer models used in this study. This research was performed with grant support from the Department of Defense Peer Reviewed Cancer Research Program CDMRP PRCRP W81XWH-21-1-0575 (to W. Gmeiner and K. Palle). This work was also supported, in part, by the Weitlauf Endowment for Cancer Research to K. Palle.

## Author Contributions

Conceived and designed the experiments: KP, NS, MBR, and WG. Performed the experiments: NS, TRO, GA, PL, HP, and CM. Analyzed the data: NS, SK, CM. Interpreted the data and contributed to the discussion: NS, SK, NC, MBR, and KP. Contributed reagents/materials/resources: KP, WG, NC, and MBR. Wrote the paper: NS and KP. Reviewed and edited the manuscript: NS, TRO, SK, GA, HP, PL, CM, WG, NC, MBR, and KP. Acquired funding: KP and WG.

