## Supplementary Macro S1 for "Novel polymeric fluoropyrimidine CF10 demonstrates superior therapeutic index and survival advantage in patient-derived models of 5-fluorouracil-refractory colorectal cancer"

**Fiji/ImageJ script for automated organoid segmentation and area quantification.**

// ---- Tune these if needed ----

rollingBall = 100; // background subtraction

threshMethod = "Li"; // "Li" is conservative; try "Triangle" if too strict

minAreaPx2 = 500; // removes tiny debris (VERY important)

excludeEdges = true; // set false if you want edge objects counted

micronPerPixel = 1.25;

pixelArea_um2 = micronPerPixel * micronPerPixel;

top = getDirectory("Choose TOP folder (includes subfolders)");

if (top=="") exit("No folder chosen.");

outCSV = top + "Holistic_Area_Recursive.csv";

if (File.exists(outCSV)) File.delete(outCSV);

File.append("Image,Width,Height,MaskPixels,Area_px2,Area_um2,PercentArea\n", outCSV);

setBatchMode(true);

files = newArray();

files = listFilesRecursive(top, files);

for (i=0; i<files.length; i++) {

path = files[i];

low = toLowerCase(path);

if (!endsWith(low, ".tif") && !endsWith(low, ".tiff")) continue;

// Open (OME-TIFF safe)

run("Bio-Formats Importer", "open=[" + path + "] autoscale color_mode=Default view=Hyperstack stack_order=XYCZT quiet");

// Force plain 2D (avoid stack dialogs)

origTitle = getTitle();

run("Duplicate...", "title=Work duplicate");

dupTitle = getTitle();

selectWindow(origTitle); close();

selectWindow(dupTitle);

run("8-bit");

run("Subtract Background...", "rolling=" + rollingBall + " sliding");

// Dark organoids on white background

setAutoThreshold(threshMethod + " dark");

// Make mask from threshold

run("Create Mask"); // creates a new mask window

maskTitle = getTitle();

// Auto-invert if background got selected (mask mostly white)

getDimensions(w, h, c, z, t);

totalPix = w * h;

hist = newArray(256);

getRawStatistics(nPix, mean, min, max, std, hist);

whitePct = (hist[255] / totalPix) * 100;

if (whitePct > 60) run("Invert"); // if >60% white, likely selected background

// Remove tiny debris; keep organoids only

opts = "size=" + minAreaPx2 + "-Infinity show=Masks clear";

if (excludeEdges) opts = opts + " exclude";

run("Analyze Particles...", opts);

// Now the active window is the cleaned particle mask (white=kept organoids)

getDimensions(w2, h2, c2, z2, t2);

totalPix2 = w2 * h2;

hist2 = newArray(256);

getRawStatistics(nPix2, mean2, min2, max2, std2, hist2);

fgPix = hist2[255];

area_px2 = fgPix;

area_um2 = fgPix * pixelArea_um2;

pct = (fgPix / totalPix2) * 100;

nameOnly = getFileNameOnly(path);

File.append(nameOnly + "," + w2 + "," + h2 + "," + fgPix + "," + area_px2 + "," + area_um2 + "," + d2s(pct,3) + "\n", outCSV);

// Close all open images from this iteration

close(); // cleaned mask

selectWindow(maskTitle); close(); // original mask

selectWindow(dupTitle); close(); // duplicated grayscale

}

setBatchMode(false);

print("DONE. Saved: " + outCSV);

}

function listFilesRecursive(dir, arr) {

list = getFileList(dir);

for (k=0; k<list.length; k++) {

p = dir + list[k];

if (File.isDirectory(p)) arr = listFilesRecursive(p, arr);

else arr = Array.concat(arr, newArray(p));

}

return arr;

}

function getFileNameOnly(fullpath) {

s = fullpath;

i = lastIndexOf(s, "\\");

j = lastIndexOf(s, "/");

k = i;

if (j > k) k = j;

if (k == -1) return s;

return substring(s, k+1);

}
